# NGN2 Expression and Regional Patterning Allow Rapid Differentiation from hiPSCs to DRG-Like Neurons Responsive to Type 2 Cytokines

**DOI:** 10.1101/2025.08.27.672624

**Authors:** Carina Habich, Loan N Miller, Nivea Granillo Luz, Laraib Iqbal Malik, Kavya Chegireddy, Mohammad Mehdi Maneshi, Michael Wolf, Markus Kummer, Hisato Iriki, Julian Röwe, Samuele Marro, Victoria E Scott, Kathleen M Smith, Brian S. Kim, Peter Reinhardt, Eric R Goedken

## Abstract

Itch or pruritus, is a sensation that elicits scratching behaviour and is a major symptom and cause of morbidity in skin diseases such as atopic dermatitis (AD), allergic contact dermatitis (ACD), prurigo nodularis (PN), and urticaria. Itch is often triggered by inflammatory stimuli in the skin including type 2 cytokines such as IL-4, IL-13, and/or IL-31. Several therapies targeting type 2 immune pathways have been developed to treat pruritus; however, itch improvement in many patients remains to be improved. Thus, additional approaches to modulate sensory neuron activity are needed. *Ex* vivo or even *in vitro* study of the molecular mechanisms underlying primary sensory neuron activation is challenging since harvesting neurons from dorsal root ganglia (DRG) in patients can only be done from cadavers. Herein, we describe rapid human sensory neurons generation (2 days to precursor cells) by *in vitro* differentiation of human induced pluripotent stem cells (hiPSC) from simultaneous application of patterning factors with *NGN2* overexpression. We show that these hiPSC-derived sensory neurons possess key characteristics of *primary* sensory neurons. They express key neuronal markers, such as TRKA receptors, TRPV1 and TRPA1 channels, and functionally respond to the TRPV1 agonist capsaicin. In addition, they express key type 2 cytokine receptors such as interleukin (IL)-4Rα and IL31-Rα, known to promote itch in AD and PN. Moreover, these cells are functional as our sensory neurons respond to IL-4, IL-13 and IL-31 stimulation. Collectively, these data demonstrate that our protocol generates a phenotypic profile consistent with native somatosensory neurons that can facilitate development of novel approaches to model and treat pruritic disease.

## Introduction

Pruritus is the central symptom in inflammatory skin diseases like AD and PN [63,64], resulting from neuroimmune interactions involving peripheral somatosensory neurons whose cell bodies are housed in the dorsal root ganglion (DRG); a subset of these neurons, or pruriceptors, detect transmit itch signals to the spinal cord and brain to initiative scratching behavior [35,42,66].

Type 2 immune cells play a pivotal role in mediating type 2 immune responses via the production of cytokines such as IL-4, IL-13, and IL-31. These cytokines contribute to inflammatory skin conditions by skin barrier disruption, promoting sensory nerve growth, and promoting neuroimmune interactions, which collectively exacerbate chronic itch and inflammation in AD and other pruritic conditions [29,57]. In AD, Janus kinase 1 (JAK1) and upstream receptor components like IL-4Rα and IL31Rα, expressed on DRG sensory neurons, are crucial for modulating immune responses and neuronal excitability related to itch [7] . These JAK1-dependent cytokine receptors are found on DRG neurons that also express canonical ion channels like TRPV1. An important aspect of pruritus stems from the effects of IL-31 from type 2 immune cells to activate pruriceptors [7]. Additionally, IL-4 and IL-13 cytokines drive type 2 inflammation impair, skin barrier, and enhance the capacity of sensory neurons to promote pruritus through TRP channels [26,36,49]. Approved treatments for AD include JAK1-selective inhibitors and antibodies neutralizing the effects of these key cytokines [14,52].

Recent single cell RNA-seq studies on primary human DRG neurons have demonstrated that somatosensory neurons are readily identified based on transcripts for various neutrophin receptors and ion channels [55]. Strikingly, pruriceptors are readily identified based on the expression of cytokine receptors such as *IL31RA*, neuropeptides such as *NPPB*, and *JAK1*. However, studying primary somatosensory neurons is challenging given that they can only be harvested from cadaveric human donors.

Human induced pluripotent stem cells (hiPSC) are a promising tool for generating expansive populations of somatosensory neurons. Differentiation protocols involve modulating signalling factors and regional patterning, beginning with neuroectodermal formation through dual SMAD inhibition of BMP/TGFβ signalling, often involving WNT signalling to specify neural crest and peripheral/sensory neurons [9,19,24]. Posteriorization in development is caused by WNT and retinoid signalling [5]. Some protocols use low cell density during neural induction [8], leading to neural crest formation [55,64]. DRG-like neurons express more posterior genes like *homeobox* (*HOX)* genes [43]. A different approach uses neuromesodermal progenitors induced by WNT, FGF signalling for posteriorization [40], sometimes combined with dual SMAD inhibition [72], which can further specify into peripheral nervous system (PNS) progeny cells. *NGN2* overexpression is a popular technique for rapidly generating hiPSC-derived neurons [11,30,73,76], although it does not yield a single neuron type [30], and some neurons express telencephalic/cortical markers like *CUX1/2* [73], but lack *FOXG1* and also induces PNS neuron markers [11,31,39]. Notably, most differentiation protocols are time-consuming and fail to yield a precise neuron population essential for itch-modeling [9,27,28,67,75].

Herein we have developed the induced DRG (iDRG) protocol for differentiating DRG neurons, combining *NGN2* induction with WNT and RA signalling. iDRG neurons express DRG markers like Trk-receptors and respond to key stimuli. They also express cytokine-signalling receptors such as IL31-Rα & IL4-Rα. Notably, JAK1 inhibition prevents cytokine activation of these cells, enabling *in vitro* exploration of the itch pathways in critical diseases like AD and PN.

## Results

### NGN2 overexpression in hiPSCs with additional patterning results in differentiation of neurons expressing markers of the dorsal root ganglia

Starting from published protocols employing *NGN2* overexpression (“iNGN2”) [46,65,76], we took advantage of two additional patterning factors, CHIR99021 (“CHIR”) & retinoic acid (“RA”), to drive sensory neuron differentiation from iPSCs (Figure 1A). In this differentiation protocol we applied these at the same time as the addition of doxycycline that induces the overexpression of the helix-loop-helix transcription factor NGN2 by an NGN2 cassette (Figure S1A). CHIR, a GSK3 inhibitor and canonical WNT activator, promotes posteriorization, dorsalization [10] and differentiation toward the neural crest whereas RA promotes posteriorization towards the spinal cord / somite region [5,22]. We found that 2 days of treatment of hiPSC with these neural induction reagents leads to differentiation of neural progenitor cells (NPC) (Figure 1B), which further differentiate into mature neurons in ∼14 days. Interestingly, it is only with the synergistic effects from both WNT and RA signalling that we observe strong expression of posterior *HOX* genes and attenuate expression of the more anterior *HOX* genes when compared to the addition of either morphogen alone (Figure S1B/C). Notably, these cells can be cryopreserved with high viability of 84.15±3.05% after thawing NPCs (e.g. for screening experiments or repetitively performed assays) on day 2, (when replating the cells), which adds to the versatility and convenience of this protocol. Moreover, due to the proliferative nature of hiPSCs, our protocol can be easily expanded to allow for the cryopreservation of large batches of NPCs. Both of these attributes of the iDRG cells enable flexibility and expanded throughput for screening experiments in plate-based assays. After two weeks of maturation, the iDRG neurons showed high expression levels of somatosensory genes like *PRDM12*, *POU4F1* (BRN3A), *ISL1*, *SCN9A* (NAV7.1), *PRPH*, *CALCA*, *CALCB* and *NTRK1* (Figure 1C). Furthermore, we also noted elevated expression of the itch-related receptors *TRPV1* and *TRPA1* (note: *TRPA1* is expressed in hiPSC [27] ) [13]. We also observed significant expression of the mature neuron marker MAPT as well as β3-tubulin expression via immunofluorescence (IF) staining (Figure 1D). Additional IF staining revealed high expression levels of TRPV1, TRPA1 and TRKA. Notably, we found there was no benefit of patterning neurons with higher concentrations of CHIR/RA (Figure S1D).

**Figure 1:**
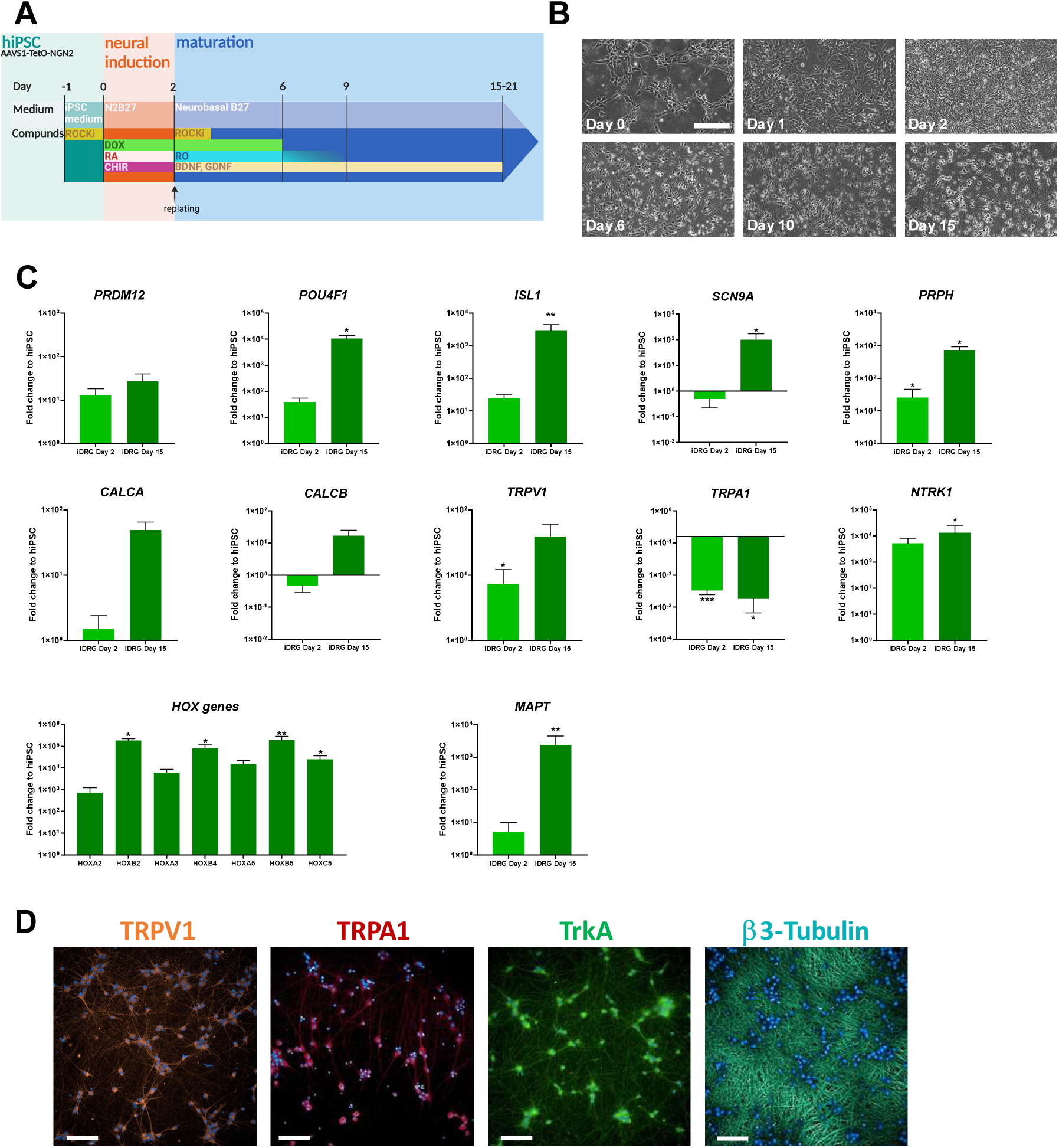
**A** Schematic representation of iDRG protocol. **B** Images of different maturation stages of iDRG (N=1 (hiPSC_1), scale bar: 200 µm). **C** q-RT-PCR examined expression of sensory neuron marker genes. iDRG neurons on day 2 (NPC) and 14 days after final replating were evaluated (N=2 (hiPSC_1, hiPSC_2), n=4, means ±SEM, * = p < 0.05, ** = p < 0.01, *** = p < 0.001). **D** Representative immunofluorescence pictures of two weeks matured iDRG neurons. A high expression of TRPV1, TRPA1, TrkA and β3-Tubulin can be shown (hiPSC_1, scale bar: 100 µm).

### Addition of neurotrophic factors after neural induction leads to a mixture of sensory neuron subpopulations responsive to itch

To increase the functionality of the iDRG, we tested if addition of neurotrophic factors could promote survival, differentiation and increase synaptic plasticity during the “maturation” stage of neurons (i.e. after the final plating and culturing time to achieve neuronal functionality and maturity). We assessed the addition of three neurotrophins (NGF (nerve growth factor), NT-3, NT-4) to the Day 2-21 (maturation) medium of iDRGs to determine their effects on the neurons’ differentiation state, as well as on their somatosensoric neuron subtype specification. Since neural precursor cells emerged after only two days of NGN2 expression, we hypothesized that these early cells would retain sensitivity to patterning factors.

NGF has a high affinity for the receptor tyrosine kinase A (TRKA, gene: *NTRK1*), whereas neurotrophin 4 (NT-4) and brain derived neurotrophic factor (BDNF) have increased binding affinities for receptor tyrosine kinase B (TRKB, gene: *NTRK2*). neurotrophin 3 (NT-3) binds to receptor tyrosine kinase C (TRKC, gene: *NTRK3*). Our iDRG switch NT/NGF condition shows the highest expression of *NTRK1*, *NTRK2* and *NTRK3*, which can be activated by the addition of NT-3, NT-4 and NGF. These genes encode receptors responding to the patterning factors we used. NGF has a high affinity for the receptor tyrosine kinase A (TRKA, gene: *NTRK1*). Neurotrophin 4 (NT-4) and brain derived neurotrophic factor (BDNF) binds more tightly to receptor tyrosine kinase B (TRKB, gene: *NTRK2*) whereas neurotrophin 3 (NT-3) activates to receptor tyrosine kinase C (TRKC, gene: *NTRK3*). Importantly stimulating receptor tyrosine kinases can trigger signalling leading to the activation of phosphorytidylinositol-3-kinase and TRPV1 [4] and NT-4 directly influences the sensitivity of nociceptors [61]. Furthermore, following addition of NGF, these nociceptors have been reported to exhibit a 20-fold higher TRKA expression [18].

We therefore carried out two treatments with NT/NGF (NT-3, NT-4, NGF) (Figure 2A). To test how NT/NGF affects maturing neurons, they were replated directly into Day 2-21 maturation medium containing NT/NGF (“iDRG NT/NGF”). To allow cells to begin to differentiate and then be exposed to the neurotrophins, the NPC were replated first in maturation medium and subsequently switched after one week into new media of the same composition plus NT/NGF (“iDRG switch NT/NGF”). The resulting cultures were assessed using RNA Sequencing for genome – wide gene expression analysis.

**Figure 2:**
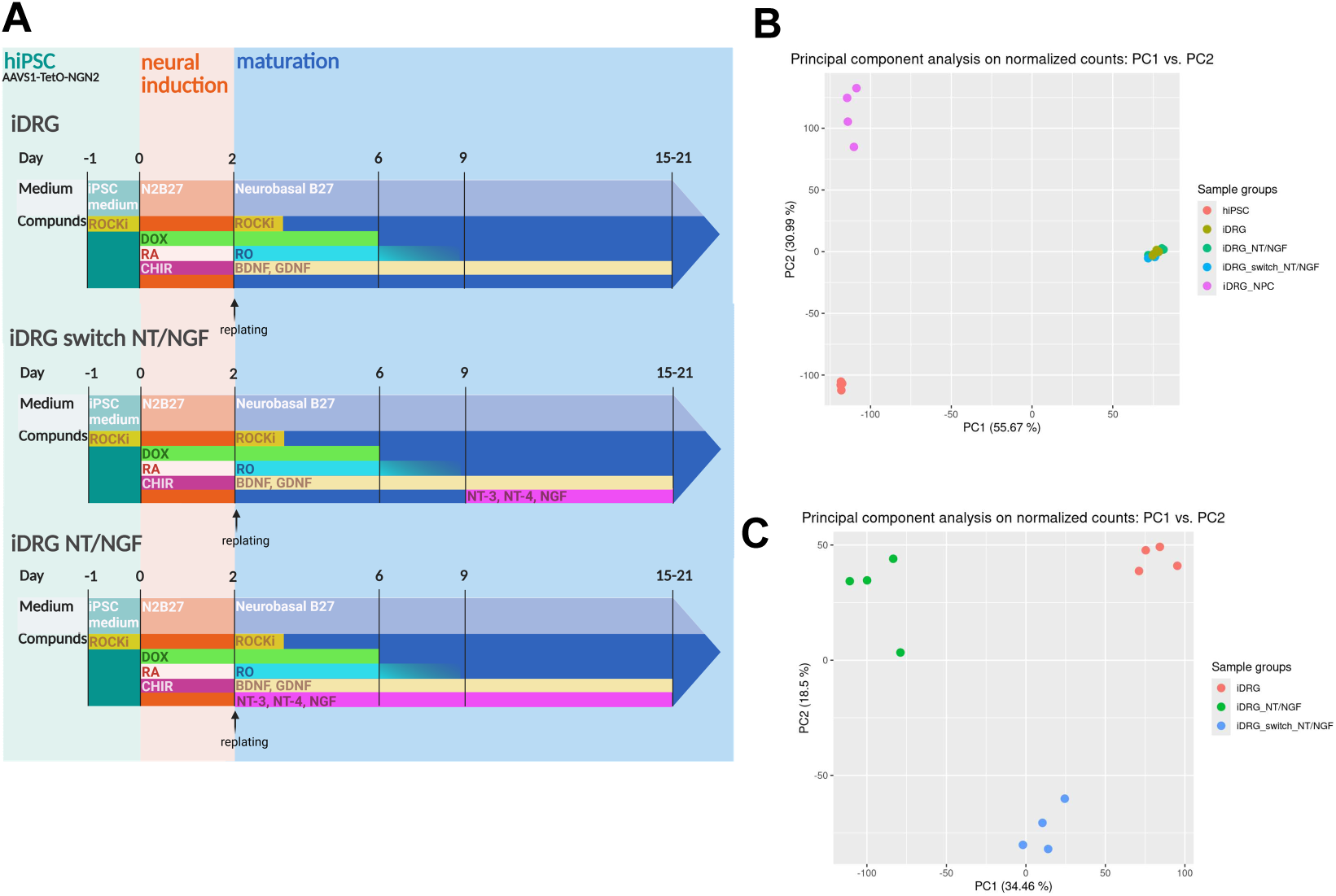
**A** Schematic representation of iDRG with different maturation conditions. **B** hiPSC were differentiated according to the protocol samples on day 2 (NPC stage) and 21 days after final replating were analysed using bulk RNA-seq. Shown is a principal component analysis (PCA) of all samples (N=1 (hiPSC_1) n=4). **C** PCA of all matured neuron conditions (N=1 (hiPSC_1) n=4).

Principal component analyses (PCA) revealed that the exclusive addition of NT-3, NT-4 and NGF affects the differentiation of iDRG neurons (Figure 2B/C) on a genome-wide expression level. Comparing hiPSC with day 2 iDRG NPC, we found an increased expression of sensory and mature neuron genes (Figure S2A). We also noted a robust induction of somatosensory neuron-like subtype mixture after the addition of NT-3, NT-4 and NGF (Figure S2B/C/D). Interestingly, there is a strong difference in gene expression between the Day 2-21 maturation conditions, differing only in the timepoint for the addition of these neurotrophins (Figure 2C). Notably, all three conditions tested show high expression levels of mature neuron marker genes (*MAPT, MAP2, TUBB3*) and marker genes associated with sensory neurons (*SST, OSMR, ISL1, POU4F1*) (Figure 3A). The expression of *NTRK1* (TRKA) and *NTRK2* (TRKB) also increases in iDRG NT/NGF and iDRG switch NT/NGF. We note that the expression of *NTRK3* (TRKC) decreases (Figure 3A/B), similar to previous reports showing that the expression of *NTRK3* decreases during maturation of the DRG [17]. Interestingly, proprioceptive genes are only weakly expressed in all neurons. Mechanosensitive related genes are expressed highest in iDRG neurons, whereas nociceptive genes (*PRDM12, P2RX3*) are significantly elevated relative to starting hiPSC cellsin iDRG switch NT/NGF neurons (Figure 3A/B). In general, none of the three conditions tested expressed more telencephalic/cortical marker genes (Figure 3A). We found in the mature iDRG, iDRG switch NT/NGF and in the iDRG NT/NGF neurons that high expression levels of *NGN1* could be detected, whereas only a low expression of *NGN1* was observed in NPC (Figure S2E).

**Figure 3:**
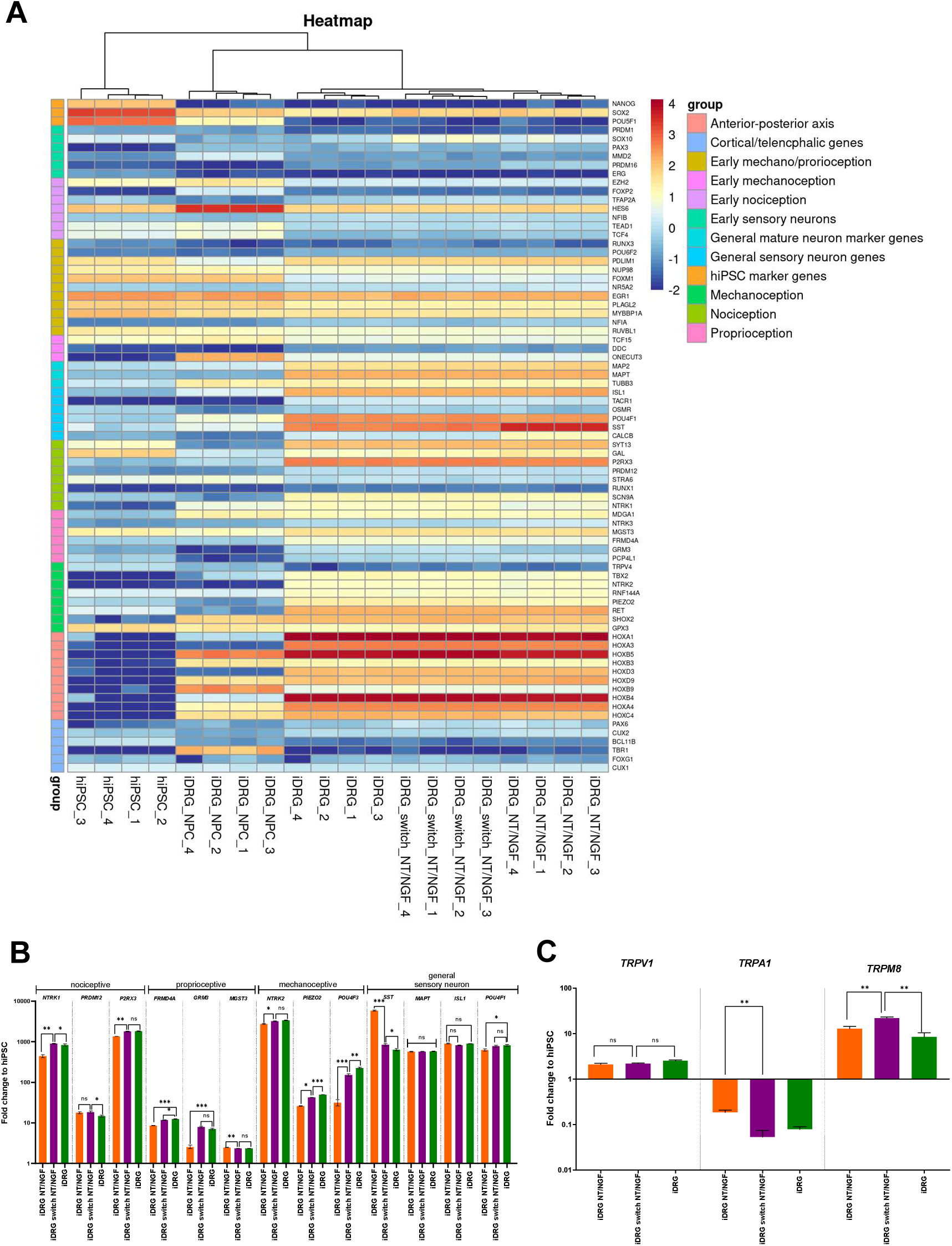
**A** Bulk RNA-seq results shown in a heatmap (N=1 (hiPSC_1) n=4). Expression levels of selected genes of interest are shown color-coded. **B** Excerpt of bulk RNA-seq data (N=1 (hiPSC_1) n=4). Fold Change of TPM against hiPSC is plotted. **C** Gene expression of relevant receptos. Excerpt of bulk RNA-seq data (N=1 (hiPSC_1) n=4). Fold Change of TPM against hiPSC is plotted.

To obtain more precise insight for future modeling itch and assessment of factors related to pathogenesis of AD, we compared expression of neuronal receptors associated with itch [13] (Figure 3C). In this analysis, the expression of *TRPV1* is similar in all conditions. Although decreased compared to the starting hiPSC, we found expression of *TRPA1* was highest in the iDRG NT/NGF subtype. *TRPM8* was expressed at highest levels in the iDRG NT/NGF switch subtype.

To test the responsiveness and electrical functionality of the various matured neurons, a multi electrode array (MEA) assay was performed. To achieve the optimal balance between maturity and reduced culture time for functionality assessments, we explored neuronal activity over the duration of neuron maturation. We found that a high mean firing of 0.55 Hz and a burst frequency of around 20 bursts per minute were achieved after about 3 weeks of maturation (day 22), with a further increase in the mean firing rate and burst frequency observed at day 25. To produce neurons with sufficient activity during the measurement while retaining the ability to increase their mean firing rate, we decided to carry out the measurements at 3 weeks after final replating (Figure 4A).

**Figure 4:**
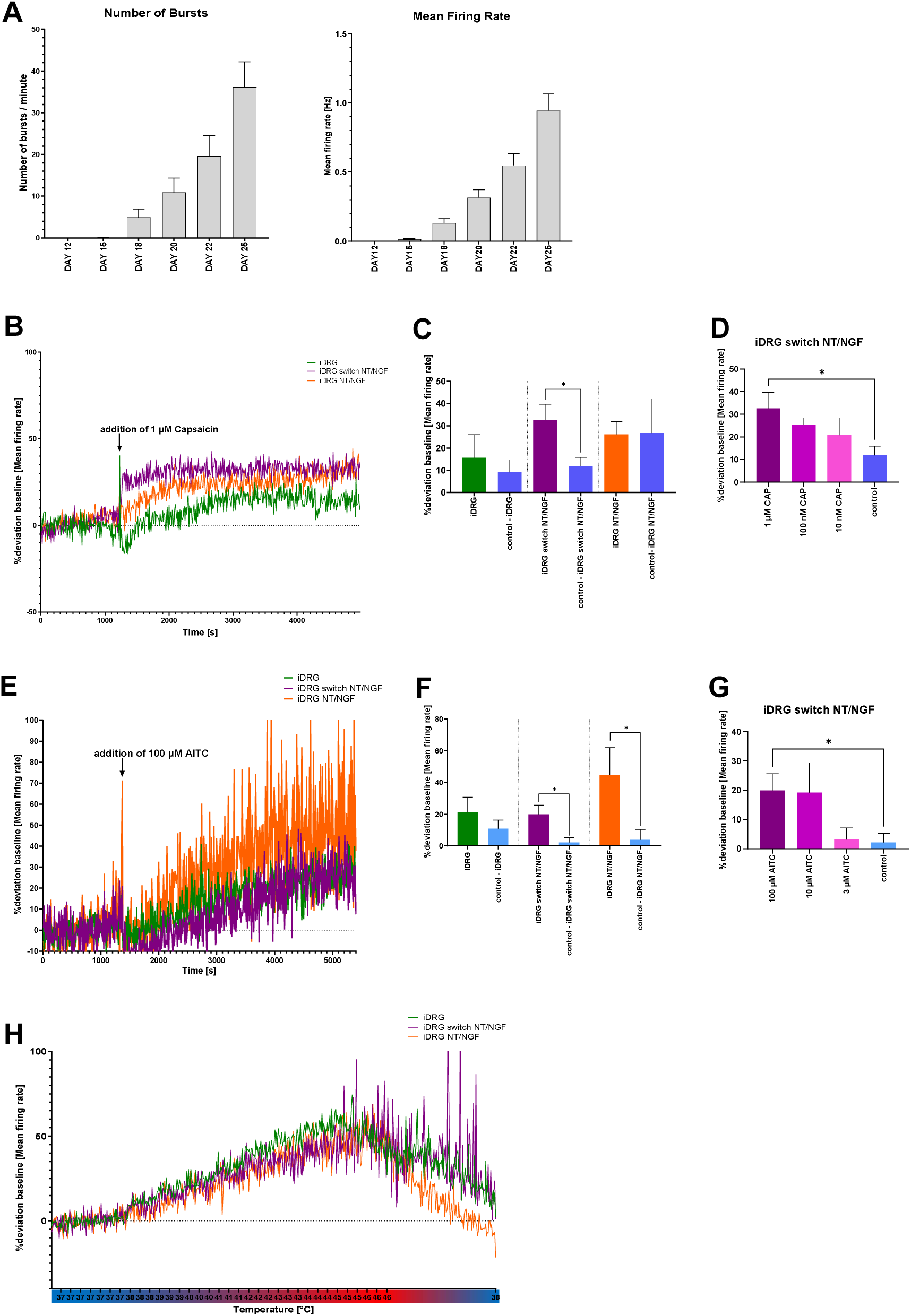
**A** MEA measurement of spontaneous activity of iDRG neurons over 25 days of maturation. Number of busts/minute and mean firing rate [Hz] are plotted (N=2 (hiPSC_1,2), n=24). **B** Addition of 1 µM capsaicin to 21 days matured iDRG, iDRG NT/NGF and iDRG switch NT/NGF neurons. A 30-minute baseline measurement was used as a zero value and the respective deviation of each well from the baseline was calculated (N=2 (hiPSC_1,2), n=11/8/12). **C** Evaluation of 1 µM capsaicin addition (Mean of 30 min after capsaicin addition ±SEM, N=2 cell lines, n=11/6/8/7/12/6). **D** Deviation of baseline after addition of different concentrations capsaicin to 21 days matured iDRG switch NT/NGF (Mean of 30 min after capsaicin addition ±SEM, N=2 (hiPSC_1,2), n=9/8/12/7). **E** Measurement via MEA of spontaneous activity after addition of 100 µM AITC (mustard oil). A 30-minute baseline measurement was used as a zero value and the respective deviation of each well from the baseline was calculated (N=2 (hiPSC_1,2), n=9/8/12). **F** Evaluation of 100 µM AITC addition (Mean of 30 min after AITC addition ±SEM, N=2 cell lines, n=9/8/8/6/12/5). **G** Measurement of activity of iDRG switch neurons (21 days matured) to addition of different concentrations of AITC (Mean of 30 min after AITC addition ±SEM, N=2 (hiPSC_1,2), n=8/8/7/8). **H** Measurement of 21 days matured iDRG, iDRG switch NT/NGF and iDRG NT/NGF spontaneous activity during increasing and decreasing of temperature (N=2 (hiPSC_1,2), n=11/11/11).

After the addition of capsaicin, we detected a significantly increased mean firing rate (32.6% compared to baseline) by TRPV1 activation in the iDRG switch NT/NGF subtype (Figure 4B/C). Notably, these capsaicin effects on iDRG switch NT/NGF were concentration-dependent (Figure 4D). In contrast, the iDRG NT/NGF neurons showed a more modest increase in mean firing rate (26.3%) and the iDRG subtype only to 15.7%. This was despite the observation that in all three differentiation conditions tested, the expression of the capsaicin-responsive TRPV1 channel is similarly elevated (Figure 3C). Thus, while all three conditions respond to capsaicin, the increased differentiated nociceptive population of the iDRG switch NT/NGF appears to drive the strongest response (Figure 4B/C).

We also measured the activity of TRPA1 in our neurons. For this purpose, allyl-isothiocyanate (AITC), the active substance in mustard oil, was added during the MEA measurement. As expected, iDRG NT/NGF showed the highest response to AITC (Figure 4E/F), since *TRPA1* is most strongly expressed in these neurons. However, like the iDRG NT/NGF subtype, iDRG switch NT/NGF also showed a significant response to AITC (Figure 4E/F), which was concentration-dependent (Figure 4G). The effects of capsaicin and AITC on iDRG switch NT/NGF neurons could also be reproduced by Ca^2+^-imaging (Figure 43A).

We next explored changes in temperature (Figure 4H). At first with early changes in temperature, all maturation conditions reacted similarly to the temperature increases. However, we found that at ∼ 45°C, there were increased effects in iDRG switch NT/NGF neurons. This was expected since iDRG switch also showed the highest response to capsaicin, and TRPV1 can also be activated with noxious heat >43°C. However, upon subsequent cooling, an increased activity of iDRG switch NT/NGF remained evident. This may reflect either a delayed reaction of TRPV1 channels or perhaps an activation of TRPM8, a receptor expressed significantly higher in iDRG switch NT/NGF than the other two conditions and is also known to be activated at moderate heat regimes and may have been triggered by the temperature changes.

After these early functional experiments, we selected the current mixture of neurons expressed in iDRG switch NT/NGF as our preferred condition to model itch. This was both because this condition responded well to the itch related receptors TRPV1 and TRPA1 [6,13] and it showed the highest increased expression of nociceptive marker genes and itch-related receptors.

### Modeling itch *in vitro* using iDRG neurons

The most relevant interleukins to modulating itch/pruritis in sensory neurons are IL-4, IL-13 and IL-31. Expression analysis of the relevant interleukin receptors and of the tyrosine kinase JAK1, which is involved in signalling of these cytokines [16,41] revealed that they were significantly increased in the matured iDRG switch NT/NGF neurons compared to the NPC (Figure 5A). When these receptors are activated by adding the corresponding interleukins to the neurons in our multielectrode array (MEA) assays, subsequent functional responses of these receptors were observed. In line with the relative expression levels of the interleukin receptors (Figure 5B), IL-4 (6.5%) and IL-13 (8.5%) induced smaller increases in the mean firing rate compared to IL-31 (19.6%) (Figure 5B). Enhanced activity over the spontaneous neuron activity after addition of IL-4, IL-13 and IL-31 could also be shown by assessing neuronal function and activity via Ca^2+^-imaging (Figure S3B/C).

**Figure 5:**
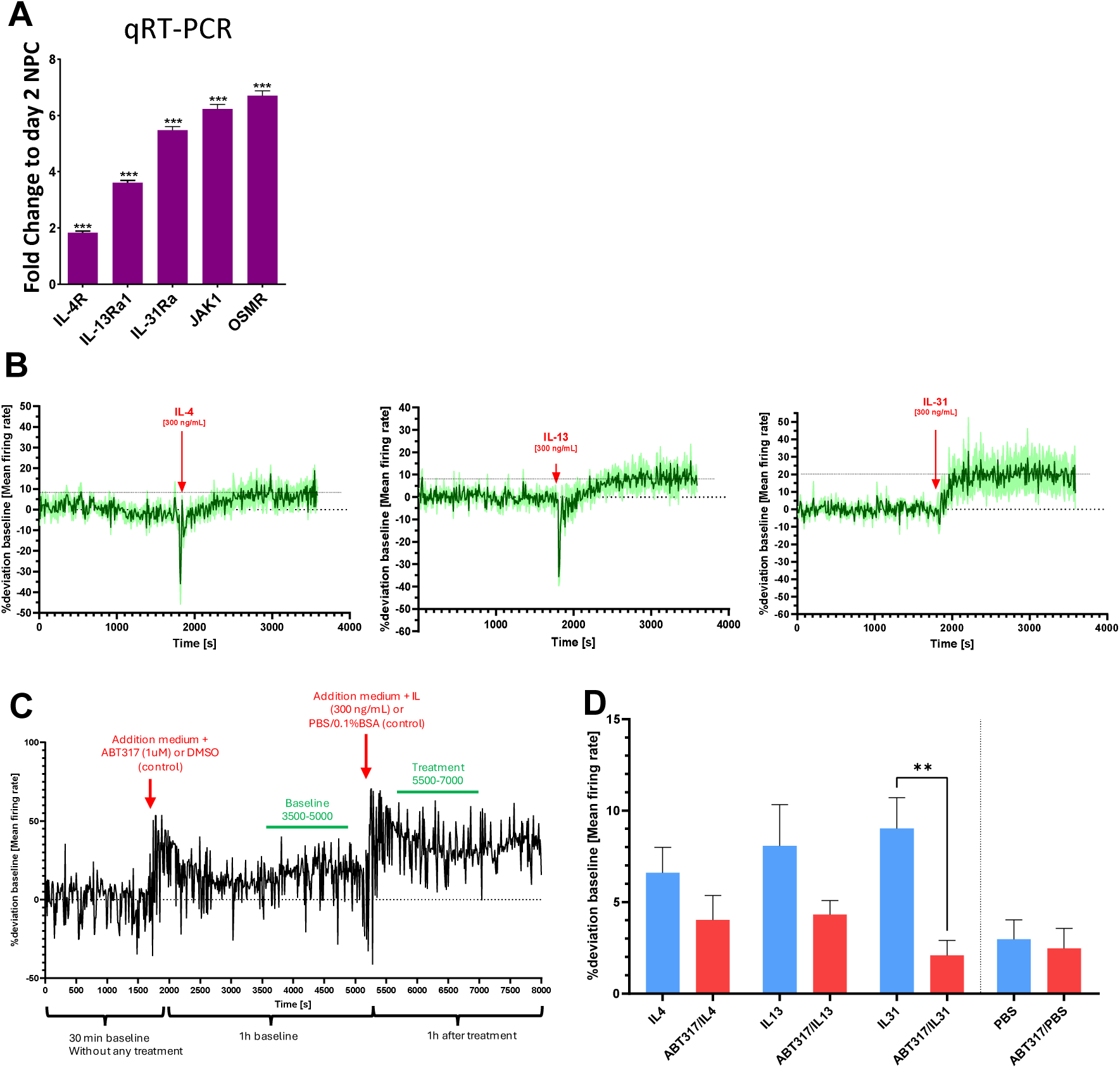
**A** qRT PCR expression of itch relevant receptors/genes of 3 weeks matured iDRG switch NT/NGF neurons. The fold change against day 2 NPC is shown (hiPSC_1, n=4, mean±SEM). **B** Reaction of 21 day matured iDRG switch NT/NGF neurons to 300 ng/mL IL-4, IL-13 and IL-31 (Light green background shows ±SEM, dark green line mean of n=4/3/3, hiPSC_1). **C** Representation of measurement time course of the effects of IL-4, IL-13 and IL-31 and JAK1inhibition (N=1 (hiPSC_1), n=1). **D** Effects of IL-4, IL-13 and IL-31 and JAK1 inhibition to 21 days matured iDRG switch NT/NGF (evaluated as presented in C, baseline 3500-5000 sec after start of measurement, treatment 5500-7000 sec after start of measurement).

After demonstrating the activation of the interleukins in iDRG switch NT/NGF subtype, we tested whether the response could be pharmacologically modulated by inhibiting JAK1. For this purpose, a baseline with the spontaneous activity of the neurons was first recorded and a JAK1 inhibitor (JAK1i) was subsequently added into the medium. For this purpose, we used ABT-317, a tool compound previously utilized in preclinical studies [21,68,69]. After a preincubation period of 1 hour, the interleukins were added (Figure 5C/D). Notably, the increase in the mean firing rate was reduced by the addition of JAK1i for all interleukins compared to the solvent control, with significant inhibition achieved with IL-31. This is not surprising as IL31RA was also most strongly expressed in these neurons, and IL-31 produced the largest stimulatory response prior to addition of JAK1i.

## Discussion

To generate translatable *in vitro* models of neurological disorders and for diseases with a sensory component, pluripotent stem cell-derived neurons have become widely used and have proven to be a valuable tool. Engineering neural subtypes from human iPSCs aids in their comparison and use in modelling even inflammatory disorders such as AD and related pruritic skin diseases. Importantly, hiPSC-derived sensory neurons provide a more patient-relevant context and can overcome translational limitations with animal models for studying human pruritus mechanisms [51]. As such, these differentiated human neurons overcome both interspecies differences and sample scarcity from animal-derived DRG neurons, offering a stable and consistent platform for itch research and drug compound screening. Thus, hiPSC-derived neurons are a valuable tool for developing and testing antipruritic agents [27].

Using iPSC-derived neurons aligns with the movement towards human cell-based systems to reducing the reliance on animal models and promoting ethical research. Furthermore, this system enables advanced measurement of neuronal responses to interactions with immune and skin cells in controlled environments [51].

These key needs – the limited availability of primary cells and the need for translatable human models – have led to protocols for differentiating disease-relevant neurons from hiPSC [9,11,19,24,30,70,73,76]. These protocols fall into two main categories: those using signalling molecules (inhibitors and activators) [9,19,24,70] and those employing timed overexpression of transcription factors specific to cell types [31,58]. The first category involves inhibiting SMAD signalling pathways (TGF/Activin/Nodal and BMP) and activating WNT signalling to induce neural crest tissue. Studies have revealed that low cell density or gamma-secretase inhibitors (such as DAPT) can increase PNS/neural crest formation over CNS [8,53] . Nevertheless, these protocols often require over a week of neural induction and patterning or replating during culture. Besides inducing neuroectoderm via dSMADi, generating neuromesoderm progenitors by WNT and FGF activators has been employed, which further patterns toward neural crest derivatives that express posterior *HOX* genes, but this laborious scheme requires multiple patterning stages and replating [25].

The second category, the overexpression of proneural transcription factors, allows for the rapid generation of neuronally-committed progeny. A very prominent gene activated in this instance is *NGN2*, and the *iNGN2* paradigm is widely used for modelling neurons from a number of disease states. Various hiPSC lines, modified to include an inducible construct at a safe harbor locus, are readily available to the scientific community. Interestingly, studies indicate that *iNGN2* neurons consist of a population with both PNS/sensory neuron components [11,30,39]. The mix of PNS and CNS is thought to arise from the transcription factor patterning. Furthermore, differentiation outcomes can be highly influenced by signalling molecules [9,19,24], making it possible to generate more specified CNS and PNS progeny.

In this study, we combined the *iNGN2* principle with a brief, strong patterning scheme. Utilizing a protocol that allows final plating or cryopreservation after just two days post-hiPSC induction, we added signalling molecules to guide cells toward a DRG-like sensory neuron phenotype. NGN2 is known to be crucial for early DRG neuron development, including the initial wave of neural crest cell migration and differentiation, while NGN1 promotes later differentiation of TRKA-positive neurons [44]. However, it has also been shown that NGN1 expression is dependent on NGN2 [2]. To enhance early sensory neuron differentiation, we chose to overexpress NGN2 instead of NGN1. NGN1 expression starts naturally during differentiation, resulting in a healthy population of TRKA-positive DRG neurons, which NGN1 overexpression alone cannot achieve [45]. *NGN1* and *NGN2* levels are highest in iDRG NT/NGF neurons, indicating matured differentiation. Despite lower *NGN1*, we fund that iDRG switch NT/NGF neurons exhibit strong *NTRK1* (TRKA) expression, proving our protocol induces adequate *NGN1* for nociceptive TRKA-positive neuron differentiation. Previous protocols utilized WNT signalling for neural crest specification and RA signalling for posteriorization. Lippmann and colleagues showed that combining WNT and FGF signalling was more effective than adding RA, though they did not test the combination of WNT/RA directly [40]. We found that combining these two stimuli led to rapid, strong upregulation of posterior HOX genes, even in just two days. Our hiPSC differentiate quickly and completely into posterior DRG-like sensory neurons without additional signalling molecules when used with the *iNGN2* paradigm, expressing relevant marker genes at both mRNA and protein levels. Addition of neurotrophins in the maturation medium (day 2-21) further enhanced neuron maturation. Compared to iDRG switch NT/NGF or iDRG NT/NGF, the iDRG neurons display more early sensory neuron markers (*PRDM16, SOX10*) [2].

The short patterning/differentiation time and the highly expandable nature of hiPSC make this protocol easily scalable, allowing for the generation of large batches for screening and repetitive assays. Crucially, it improves expandability and adaptabiity compared to lengthy protocols that require regular media changes. We also note that the protocol fits within a traditional 5-day work week and is applicable to existing hiPSC clones with an inducible NGN2 cassette which is an important neuron source in the field. Interestingly, modifying the maturation conditions significantly altered the neuronal composition, supporting the protocol’s adaptability for driving toward different conditions to enhance desired neuronal subtypes. We also note the protocol does not require glial cells and lacks astrocyte marker expression (such as *GFAP* – not shown), and single non-neuronal cells are eliminated through mitotic inactivation. Thus, our differentiation conditions yield highly functional, mature neurons, suitable for neuronal studies and, due to their purity, ideal for co-culture experiments with different cell types.

In this study, we found expression of numerous markers expected on sensory neurons. The expression of TRKA, TRPV1, and TRPA1 is particularly significant, as these proteins play crucial roles in sensory perception [15,23,48]. The presence of the ion channels TRPV1 and TRPA1, which detect thermal and chemical stimuli, suggests that our iDRG neurons are poised to respond to environmental stimuli, reinforcing their sensory neuron identity. Indeed, we have also shown these neurons respond to capsaicin, a known TRPV1 agonist. We found that the TRPV1 receptor, activated by capsaicin and noxious heat (>43°C) [62] showed the highest response in the iDRG switch NT/NGF subtype. This subtype also responded strongly to both noxious heat and cooling, which we attribute to increased expression of TRPM8, a receptor responding to menthol heat (28-15°C) and cooling (<15°C) [47]. The TRPA1 receptor is triggered by to mustard oil (AITC) and generally is activated by electrophilic substances [37,50].

In addition to somatosensory neuron markers, we observed expression of type 2 cytokine-signalling receptors like IL4-Rα, IL13-Rα1 and IL31-Rα. This enables use of these neurons in studying neuroimmune pathways linked to pruritus in AD. The roles of IL-4, IL-13, and especially IL-31 in AD pathogenesis and resulting pruritis are tied to the activation of sensory neurons, a cell type are centrally involved disease mechanisms and progression [33]. IL-31 is produced by various immune cells, and its elevated levels correlate with increased itch severity [1,20]. Moreover, IL-31Rα, when activated, not only augments neuronal excitability but also impacts other immune pathways contributing to the chronic inflammatory state characteristic of AD [54].

Indeed, the interaction between type 2 cytokines, such as IL-31 and sensory neurons is crucial in AD pathophysiology, positioning IL-31 as a critical target for therapeutic interventions. Treatments that inhibit IL-31 signalling, including IL-31Rα inhibitors like nemolizumab and downstream signalling molecules such as JAK1, have demonstrated effectiveness in alleviating pruritus and improving patient outcomes in AD [32,74].

We observed that our iDRG switch NT/NGF neurons respond to recombinant human IL-31 resulting in increased neuronal activity. Furthermore, both MEA and Ca^2+^-imaging experiments demonstrated that iDRG switch NT/NGF neurons respond to IL-4, IL-13, and IL-31. These methods are frequently used to characterize DRG activity [27,56] and show complementary data. Moreover, the excitability observed after addition of IL-31 could be significantly reduced by a JAKi (ABT-317). This demonstrates the iDRG switch NT/NGF neurons respond to a relevant drug molecule in a functional model of early itch signalling.

To further enhance the physiological relevance of hiPSC-derived DRG models, integrating co-culture systems with skin-relevant cells or tissues represents an attractive potential next step [71]. The diverse sensory neuron markers expressed in our iDRG validate their classification as sensory neurons and highlight their utility in disease modelling for process including itch and chronic pain. Since we demonstrated it is possible to skew the secretory phenotype with simple adaptations in the timed addition of signalling factors, the protocol presented here can also serve as a template for modelling other sensory neuron functions *in vitro.* The presence of cytokine signalling receptors suggests further utility for research in neuroinflammatory conditions, aiding therapeutic discovery. As such, our in vitro differentiated iDRG neurons can serve as essential tools for basic and translational pruritus research.

## Material and Methods

### Cell culture

In this study, a publicly available hiPSC line with an *iNGN2* cassette were used (BIONi010-C-13, available from EBiSC www.ebisc.org. The EBiSC Bank acknowledges Bioneer A/S as the source of human induced pluripotent stem cell line BIONi010-C-13 which was generated with support from EFPIA companies and the European Union (IMI-JU), Table 1). The other hiPSC line (hiPSC_2) generated from Schöndorf et *al.* ([60] , BiomedX) with an integration of an iNGN2 cassette. Both hiPSC lines can be found in Table 1.

**Table 1:** hiPSC lines.

| Name | Referred as | Source/<br>Modification | Disease<br>genotype | Sex | Age | Reprogramming<br>Method | Modification |
| --- | --- | --- | --- | --- | --- | --- | --- |
| BIONi010-C-13 | hiPSC_1 | EBISC* | Control<br>cell line | male | 15-19 | Episomal<br>Reprogramming<br>factors: KLF4, Lin28,<br>MYC, POU5F1, shP53,<br>SOX2 | edited using CRISPR at<br>CRO (Bioneer) with a<br>DOX-inducible NGN2<br>overexpression<br>construct introduced in<br>the AAVS1 alleles (one<br>allele CAG:m2rtTA, other<br>allele TRE:NGN2) |
| BIOMEDI005-A (Coriell<br>Fibroblast<br>AG06869) | hiPSC_2 | BioMedX/<br>Bioneer | sAD | fema<br>le | 60 | RNA based<br>reprogramming [60],<br>Reprogramming<br>factors: OSKM | see hiPSC_1 |

hiPSC were cultured in E8 Flex medium (Thermo Fisher) on cell culture ware coated with Matrigel (Matrigel hESC-Qualified Matrix, LDEV-free, Corning) [46]. Matrigel was applied using DMEM/F12 (Thermo Fisher) and coated for at least 24h at 4°C. The medium was changed every 2-3 days, depending on the cell density and cells routinely split twice a week using Versene / EDTA (Lonza) as clumps, or as single cells with Accutase (Thermo Fisher) supplemented with 10 μM Y-27632 (Merck, referred to as ROCKi) for 10-15 min at 37°C. The cells were then replated on Matrigel-coated cell culture ware in E8 flex medium with 10 μM ROCKi, which was removed the following day.

### Differentiation of hiPSC derived DRG

The basic framework of the protocol used is from [46,65]. For the experiments hiPSC were seeded on Matrigel coated plates, as single cells at a density of 52,000 cells/cm^2^ in E8 flex medium supplemented with 10 μM ROCKi (day -1). On day 0, cells washed once with N2B27 medium (Table 2) to remove E8 flex medium. Cells were then supplied with 0.5 mL/cm^2^ N2B27 medium supplemented with 2 μg/mL Doxycycline hydrochloride (DOX, Merck), 3 μM CHIR99021 (CHIR, tocris) and 1 μM retinoic acid (RA, all-trans retinoic acid, Merck). The following day (day 1) the medium was completely changed using the same medium. On day 2 cultures were incubated at 37°C for 15 min with prewarmed accutase containing 10 μM ROCKi. The single cell solution was then diluted 5x in N2B27 medium and centrifuged for 5 min at 300xg. If the NPC were to be frozen, the pellet was resuspended in pre-cooled Synth-a-Freeze medium (Thermo Fisher) supplemented with 10 μM ROCKi and frozen in a CoolCell container (Thermo Fisher) at -80°C. Cryopreserved cells were transferred to liquid nitrogen for long term storage after a maximum of 3 days. In case of fresh replating, resulting pellet was resuspended in neural maturation medium (NMM, Table 3) supplemented with 2 μg/mL DOX, 10 μM ROCKi and 500 nM RO4929097 (Selleckchem). In condition “iDRG NT/NGF”, the medium was additionally supplemented with NT3 (10 ng/mL, R&D), NT4 (3 ng/mL, R&D) and NGF (30 ng/mL, R&D), “iDRG switch NT/NGF” and “iDRG” conditions were replated in medium without these additives. Cells were replated on PLO-Matrigel coated plates following the method described in Manos et al. For imaging purposes, 41,000 cells/cm2 were seeded in 0.45 mL/cm2, for RNA analysis 100,000 cells/cm^2^ in 0.42 mL/cm^2^. On day 3 a 70% medium change was performed with NMM supplemented with 2 μg/mL DOX, 5 μM ROCKi and 500 nM RO4929097 (iDRG NT/NGF with additional NT3, NT4 and NGF). Day 6, a 1 μg/mL Mitomycin C (Merck) treatment was performed for 1 h at 37°C. Following this, a complete medium change using NMM supplemented with 100 nM RO4929097 was performed. From then on, every 4-5 days, a 50% medium change with NMM was performed. In condition iDRG NT/NGF, NT3, NT4 and NGF were further added. 7 days after replating, the medium for iDRG switch NT/NGF was completely exchanged for medium supplemented with NT3, NT4 and NGF. iDRG received the complete maturation NMM without NT3, NT4 and NGF.

**Table 2:**
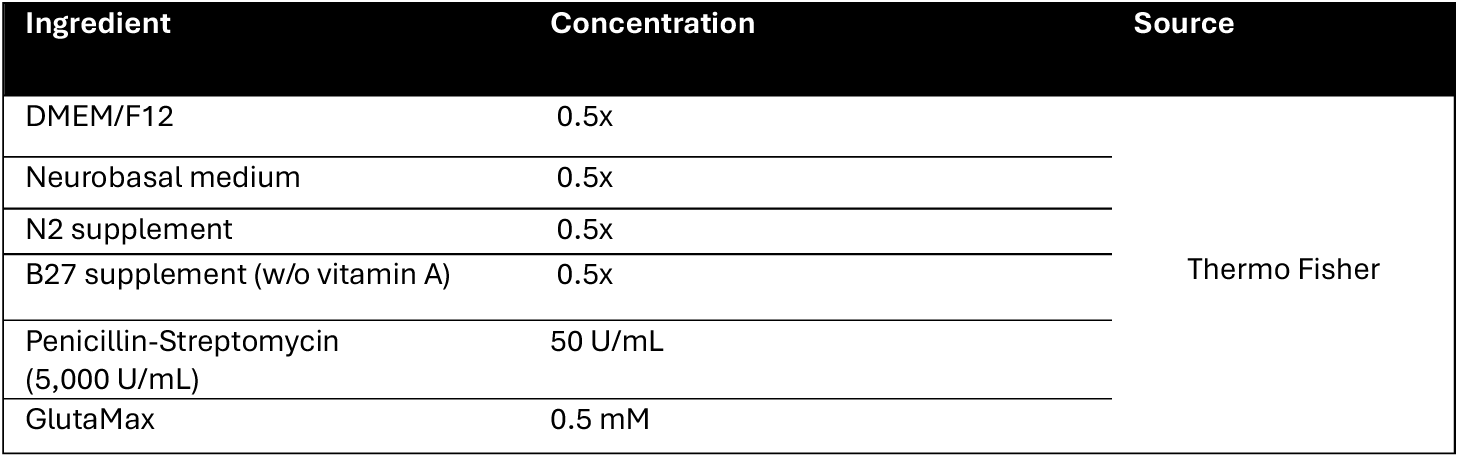
N2B27 medium.

| Ingredient | Concentration | Source |
| --- | --- | --- |
| DMEM/F12 | 0.5x | Thermo Fisher |
| Neurobasal medium | 0.5x |  |
| N2 supplement | 0.5x |  |
| B27 supplement (w/o vitamin A) | 0.5x |  |
| Penicillin-Streptomycin (5,000 U/mL) | 50 U/mL |  |
| GlutaMax | 0.5 mM |  |

**Table 3:** Neural maturation medium (NMM).

| Ingredient | Concentration | Source |
| --- | --- | --- |
| Neurobasal plus medium | 1x | Thermo Fisher |
| B27 Supplement plus supplement | 1x | Thermo Fisher |
| GlutaMax | 1 mM | Thermo Fisher |
| Penicillin-Streptomycin (5,000 U/mL) | 50 U/mL | Thermo Fisher |
| BDNF | 10 ng/mL | Biotechne |
| GDNF | 5 ng/mL | Biotechne |
| dcAMP | 200 µM | Merck |
| L-Ascorbic acid | 200 µM | Merck |
| Laminin | 2 µg/mL | Merck |

### q-RT PCR

All samples were lysed in 350 µL RLT buffer (Qiagen). RNA isolation was performed using RNeasy Plus Mini Kit (Qiagen). For cDNA synthesis, Superscript IV VILO Master Mix with ezDNase (Thermo Fisher) was used. Analysis was carried out via TaqMan Assays (Thermo Fisher, Table 4) in a 20 µL reaction using 10 ng RNA input. The qRT-PCR was done using a QuantStudio 7 Flex Real Time PCR System (Thermo Fisher) and was carried out under fast conditions (Initiation at 95°C for 20sec, 40 cycles of denaturation at 95°C for 1 sec followed by 20 sec at 60°C annealing/extension). The samples were normalized to two housekeeping genes (GAPDH, RPL13) and then calculated using 2-ΔΔCt method. The results were evaluated with Graphpad Prism 10. If CT was undetected during a probe, the CT was set to maximal number of cycles (CT=40). All CT values were normalized to the CT values of the housekeeping genes. For statistical analysis, first the normal distribution was checked via Shapiro-Wilk test. To compare two groups, an unpaired t-test was performed for equal variances and a Kolmogorov-Smirnov test for different variances. The significant differences were shown as follows: * = p < 0.05, ** = p < 0.01, *** = p < 0.001.

**Table 4:** Primer real-time PCR.

| Gene | Company | Assay ID |
| --- | --- | --- |
| GAPDH | Thermo Fisher | Hs99999905_m1 |
| CALCA | Thermo Fisher | Hs01100741_m1 |
| CALCB | Thermo Fisher | Hs00265194_m1 |
| HOXA2 | Thermo Fisher | Hs00534579_m1 |
| HOXA3 | Thermo Fisher | Hs00601076_m1 |
| HOXA5 | Thermo Fisher | Hs00430330_m1 |
| HOXB2 | Thermo Fisher | Hs01911167_s1 |
| HOXB4 | Thermo Fisher | Hs00256884_m1 |
| HOXB5 | Thermo Fisher | Hs00357820_m1 |
| HOXC5 | Thermo Fisher | Hs00232747_m1 |
| ISL1 | Thermo Fisher | Hs00158126_m1 |
| MAPT | Thermo Fisher | Hs00902192_m1 |
| NTRK1 | Thermo Fisher | Hs01021011_m1 |
| POU4F1 | Thermo Fisher | Hs00366711_m1 |
| PRDM12 | Thermo Fisher | Hs00964106_m1 |
| PRPH | Thermo Fisher | Hs00986945_g1 |
| RPL13 | Thermo Fisher | Hs00744303_s1 |
| SCN9A | Thermo Fisher | Hs01076699_m1 |
| TRPV1 | Thermo Fisher | Hs00949994_m1 |

### Immunofluorescence (IF) staining

For IF staining neurons were fixed for 15 min with 8% formaldehyde solution which was added to the culture medium to result in a 4% dilution. Afterwards, cells were washed three times with PBS (PBST, Gibco). Permeabilizing solution (0.2% TX-100 (Sigma) in PBS) was added for 10-20 min followed by blocking with 2% BSA (Invitrogen) in PBS for 30 min.

The primary antibodies (Table 5) were then diluted in antibody solution (0.6% BSA in PBS) and incubated overnight at 4°C. The plates were washed three times with PBS and then the secondary antibodies (Table 5) were diluted in antibody solution. Plates were incubated for 1 h at RT. Followed by an incubation for 10 minutes with 1:4,000 Hoechst 33342 (Invitrogen) in antibody solution. Another wash with PBS was performed. The IF pictures were taken on Thermo Fisher CX7 and the quantitative analysis of the data was carried out using the Cell Insight CX7 High Content Analysis platform (15900603). Pictures of Figure S1D were taken with Operetta CLS (Revity) and evaluated via Harmony 5.2.

**Table 5:** Primary and secondary antibodies for IF staining.

| Antibody | Supplier | Catalogue number | Dilution |
| --- | --- | --- | --- |
| Anti-MAP2 antibody | Abcam | ab5392 | 1:10,000 |
| b-3-Tubulin | Abcam | ab18207 | 1:800 |
| Chicken anti-goat Alex Flour 488 | Invitrogen | A21467 | 1:400 |
| Donkey anti-Rabbit IgG (H+L) Highly Cross-Adsorbed Secondary Antibody, Alexa Fluor Plus 488 | Thermo Fisher | A32790 | 1:1,000 |
| Goat anti- Rabbit IgG-FITC | Jackson Immunoresearch | 110-095-144 | 1:200 |
| Goat anti-Chicken IgY (H+L) Secondary Antibody, Alexa Fluor 555 | Thermo Fisher | A21437 | 1:1,000 |
| Goat anti-rabbit Alex Flour 555 | Life Tech | A21428 | 1:400 |
| Goat anti-rabbit Alex Flour 647 | Life Tech | A21428 | 1:400 |
| HOXA3 Polyclonal Antibody | Thermo Fisher | PA5-56077 | 1:100 |
| TrkA | R&D | A-F175 | 1:400 |
| TRPA1 | NOVUS | NB-110-40763 | 1:200 |
| TRPV1 | Thermo Fisher | PAI-748 | 1:400 |

#### Multi electrode array assay (MEA assay)

Using the Maestro Pro (Axion Biosystems), 30,000 cells were drop seeded onto 48 well poly(ethyleneimine) solution (PEI 50%, MERCK) pre-coated MEA plates. To coat the plates with PEI, 5 µl of freshly prepared 0.1% PEI (1 mL 10% PEI solution (2g 50% PEI stock solution, 8 mL sterile water) with 99 mL 25 mM borate buffer pH 8.4 (Merck) was added in the middle of the well and incubated at 37°C for 1 h. The wells were then washed three times with sterile water and twice with DPBS and left to dry overnight at room temperature. Next day cells were seeded in the middle of the well in MEA replate medium (7:1 NMM supplemented with 2 μg/mL DOX, 10 μM ROCKi and 500 nM RO4929097)/ 193 µg/mL Matrigel in DEMEM/F12) for 1 h at 37°C before being treated as described in the protocols.

All measurements of final neurons were done 3 weeks after final replating on Maestro Pro (Axion Biosystems) with 48 well plates. Before all measurements the MEA plates were equilibrated for 10 minutes in the Maestro Pro. For activity measurements an 8 min time slot of the plate of spontaneous firing was detected. For measurements with addition of small molecules/interleukins 1h prior to activity measurements, a 50% medium exchange was performed. All small molecules (Capsaicin, Tocris; JAK1 inhibitor (31Syntheised in-house at AbbVie); Allyl-isothiocyanat, (Merck)) and interleukins (IL-4, IL-13, IL-31 (R&D)) were dissolved as specified by manufacturer and further diluted in respective medium. The baseline was recorded for 30 min before adding first treatment. Afterwards 1h measurement of just spontaneous activity was performed. If necessary, the second treatment was then added followed by another hour of measurement of spontaneous activity. As control always the same amount of solvent (DMSO or PBS/0.1% BSA) was diluted in the respective medium as small molecules/interleukins. Measurement of IL’s and JAK1 inhibition was performed using corticosteroid free medium (NMM without corticosteroid customized, Thermo Fisher). To scale the efficiency of the inhibition, 25 minutes before addition was set as the baseline (0% value) and the increase in the baseline after the peak of the addition of the interleukins, which occurs through the mechanical stimulation of the neurons, was used as a comparison value for the stimulation. For temperature measurements first 15 min were measured as baseline, then within 1 min the temperature was raised 1°C. The temperature was held for 5 min, followed by the next step of increasing temperature. The temperature was increased to 46°C and then cooled to 38°C. The analysis was conducted using AxIS Navigator (Axion Biosystems), Neural Metric Tool (Axion Biosystems), Excel (Microsoft) and Graphpad Prism 10. For statistical analysis first a normal distribution was checked via Shapiro-Wilk test. Followed by an unpaired t-test was performed for equal variances and a Kolmogorov-Smirnov test for different variances. The significant differences were shown as follows: * = p < 0.05, ** = p < 0.01, *** = p < 0.001.

#### Calcium-imaging

Cells were plated in clear bottom black 96-well plates at a density of 35,000 cells/well, and 2 weeks matured iDRG switch NT/NGF were tested in Cystation5 (Agilent, CA). On the day of the experiment, the medium was replaced, and the cells were incubated with Fluo-4 (Invitrogen F14201) in Hanks buffered saline solution containing Ca^2+^ and Mg^2+^ (HBSS, Gibco) for 30 minutes at 37°C. The cells were then washed with HBSS buffer and rested at 37C for 30min to allow the desertification of the dye. All stimulations done acutely was done via the injectors of Cytation5 at the lowest rate of 225 µl/sec. 10 µl of the (10X) agonist was added to the well containing 90ul HBSS buffer to bring the concentration to the final concentration stated in each figure. The cells were added with fixed concentrations of capsaicin at 10 nM, 100 nM, and 1µM, respectively, as well as mustard oil at 10 µM and 100 mM respectively. Cells were imaged in widefield mode using the 4X objective and the GFP imaging cube (Ex: 469/35, EM: 525/39). Stim in each condition was added after measurement of 30 seconds baseline activity. To process the images, first background subtraction was done for each well separately. Then the sum GFP intensity of the well was calculated for each time point (F). Baseline fluorescence at t=0 (F_0_) was used to normalize the response between the wells and F/F_0_ was calculated for each well and plotted. For chronic Type 2 cytokine treatment experiments, cells were incubated for 2 hours at 37°C with recombinant human IL-4, IL-13, and IL-31 (2 µg/mL each, R&D systems). Cells were washed and labelled with Fluo-4 as described (Invitrogen F14201).

#### Bulk RNA sequencing

Bulk RNA sequencing was performed with the Illumina NExtSeq 550 System with NextSeq 500/550 High Output Kit v2.5 (Illumina, 20024906). For library preparation Illumina Stranded mRNA Prep, Ligation (20040534) and IDT for Illumina RNA UD Indexes Set A, B, C (20040553, 20040554, 20040555) were used. The denaturation of the final library was carried out according to protocol A (Denature and Dilute Libraries Guide, Illumina).The RNAseq data was trimmed for quality purposes using trim galore (version 0.6.4_dev) [34] , then aligned against human reference (GRCh38.106) using STAR (version 2.7.6a)[12] . Gene counts for each sample were collected using Rsubread package (version v2.0.0)[38] and them TMM normalized. Since the data was processed in two batches, with 4 common samples across batches, it was combined and normalized using ComBat-seq from sva (version 3.54.0)[77]. Differential gene expression analysis for pairwise comparison was done using limma (version 3.60.6)[59] . Volcano plots were generated using EnhancedVolcano (version 1.24.0)[3]. All processing was done in R (version 4.4.1).

## Acknowledgements

We would like to thank AbbVie employees Elke Käfer and Anja Fink for their technical assistance to provide frozen stocks of hiPSC. We would also like to thank Prof. Dr. Martin Grininger and Prof. Dr. Jasmin Hefendehl for their supervision. The following figures were created with BioRender.com: 1A, S1A and 2A.

## Author Contributions

Conceptualization, P.R., L.N.M, C.H. and E.R.G.; Methodology, P.R., A.K. and C.H.; Validation, C.H., L.N.M., M.M.M, K.C., K.M.S. and L.I.M.; Formal Analysis, C.H. and L.I.M., Investigation, C.H., L.N.M., M.M.M and M.W.; Data Curation, C.H., L.I.M., M.K. K.M.S. and K.C.; Writing – Original Draft, P.R., C.H., N.G.L and E.R.G.; Writing – Review & Editing, C.H., L.N.M., K.C., H.I., L.I.M, M.M.M., M.W., M.K., J.R:, S.M., V.E.S., K.M.S., B.K., P.R. and E.R.G.; Visualization, C.H., L.I.M. and L.N.M.; Supervision, P.R. and E.R.G.; Project Administration, P.R. and E.R.G..

