## Supplementary data for "NGN2 Expression and Regional Patterning Allow Rapid Differentiation from hiPSCs to DRG-Like Neurons Responsive to Type 2 Cytokines"

### Supplementary information

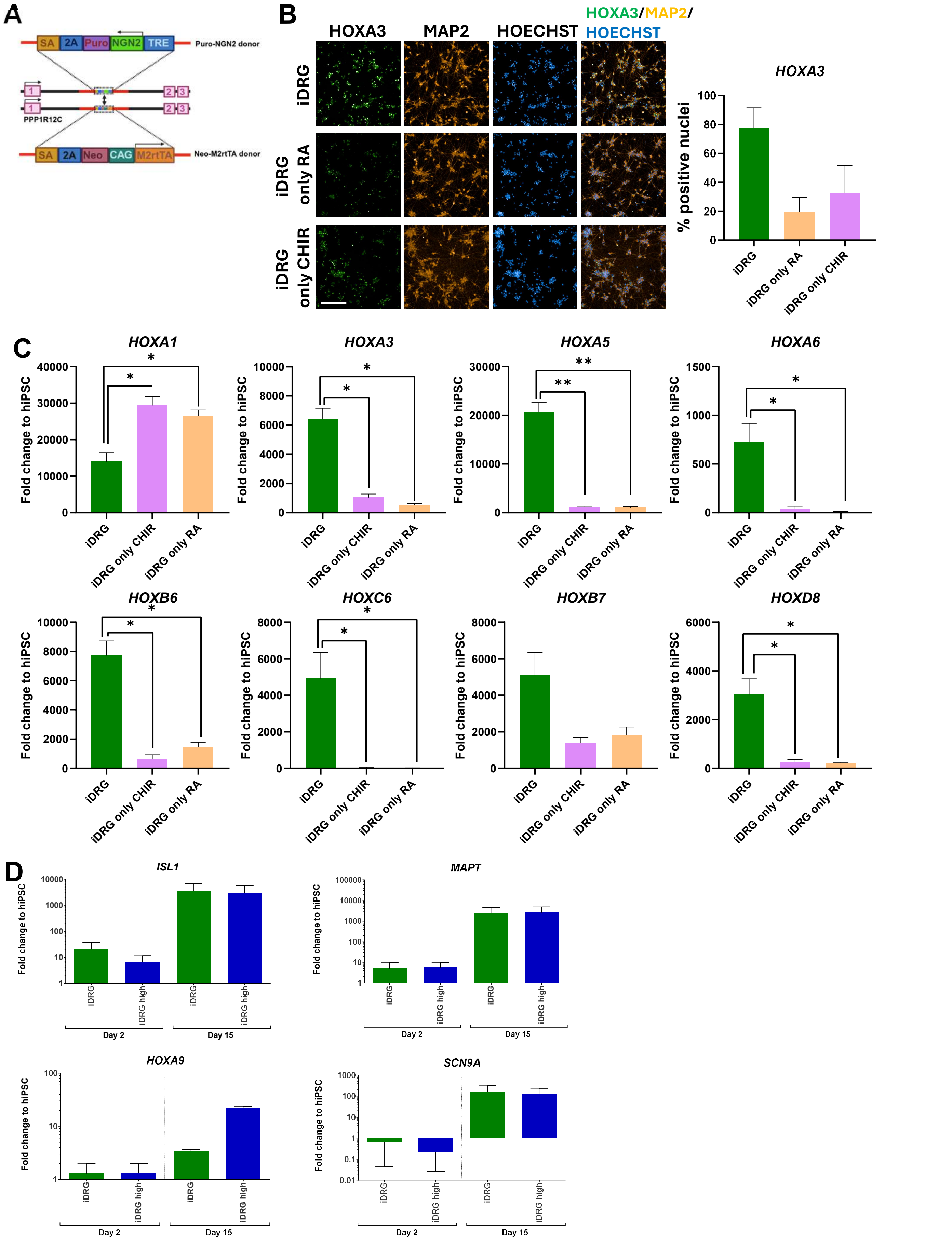


**Figure S1:** **A** Overexpression of the basic helix-loop-helix transcription factor neurogenin 2 (NGN2) facilitates the direct differentiation of human induced pluripotent stem cells (hiPSCs) into neurons. In the ASSV1 (PPP1R12C) locus, gene editing using the HDR pathway was utilized to insert puro-NGN2 and neo-M2rtTA across two alleles. The expression of NGN2 is induced by the addition of doxycycline (DOX). **B** Representative immunofluorescence pictures of two weeks matured iDRG neurons compared to “iDRG” neurons without CHIR or RA in neural induction. Quantification of HOXA3 is shown (hiPSC_1, n= 2, means ±SEM). **C** Evaluation of bulk RNA sequencing of 14 days matured iDRG neurons (hiPSC_1, n=4, scale bar: 200 µm, means ±SEM, * = p < 0.05, ** = p < 0.01, *** = p < 0.001). In neural induction part of iDRG protocol just RA or CHIR was added. These neurons were compared to iDRG neurons. **D** q-RTPCR detected expression levels of sensory marker genes. iDRG were patterned using 3 µM RA and 10 µM CHIR. After final replating the neurons were matured for two weeks (N=2 cell lines (hiPSC_1, hiPSC_2), n=4, means ±SEM, * = p < 0.05, ** = p < 0.01, *** = p < 0.001).

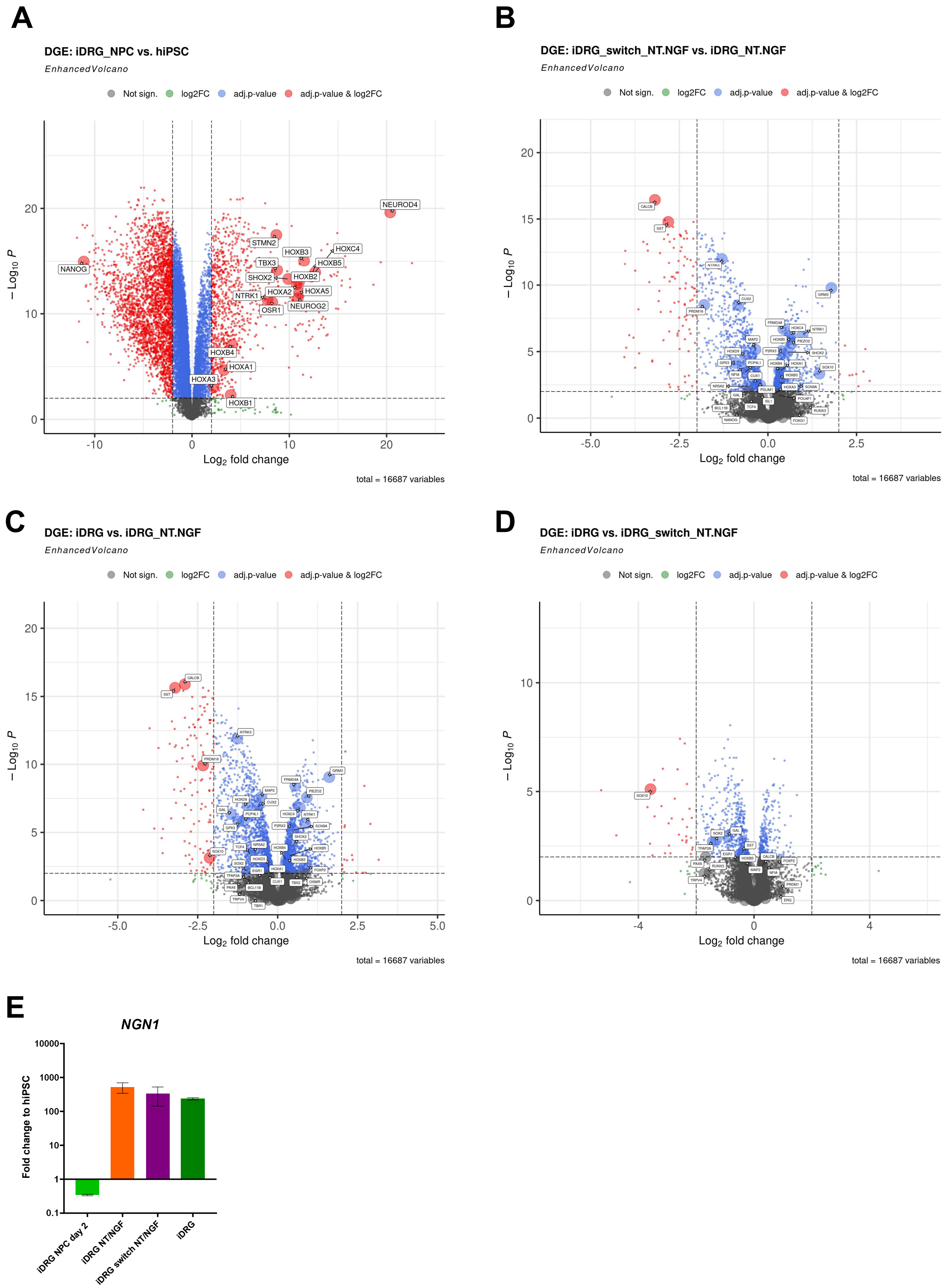


**Figure S2: A** Volcano plot of hiPSC compared with iDRG day 2 via bulk RNA-seq (data of Figure 2, N=1 (hiPSC_1), n=4). **B** Comparison of iDRG and iDRG NT/NGF neurons via volcano plot (data of Figure 2, N=1 (hiPSC_1), n=4). **C** Comparison of iDRG and iDRG switch NT/NGF neurons via volcano plot (data of Figure 2, N=1 (hiPSC_1), n=4) **D** Comparison of iDRG switch NT/NGF and iDRG NT/NGF neurons via volcano plot (data of Figure 2, N=1 (hiPSC_1), n=4). **E** Excerpt of bulk RNA-seq data (data of Figure 2, N=1 (hiPSC_1), n=4). *NGN2* and *NGN1* TPM fold changes against hiPSC are plotted.

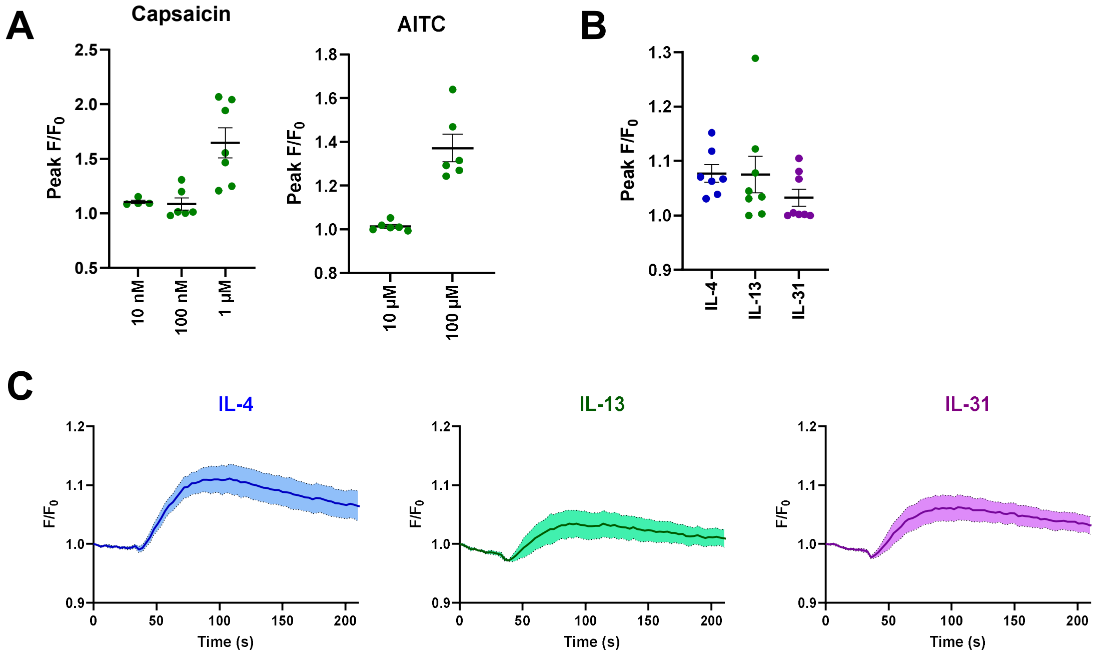


**Figure S3: A** On 14 days matured iDRG switch NT/NGF neurons $Ca^{2+}$-imaging was performed. The neurons were stimulated with CAP or AITC in different concentrations (hiPSC_1, n=6, results are shown as means ±SEM). **B** Via $Ca^{2+}$-imaging response of iDRG switch NT/NGF neuons on IL-4 (2 µg/ml), IL-13 (2 µg/ml) and IL-31 (2 µg/ml) was tested (hiPSC_1, n=7, results are shown as means ±SEM). **C** Time course of data shown in Figure S3A.
